# Magnetic voluntary head-fixation in transgenic rats enables lifetime imaging of hippocampal neurons

**DOI:** 10.1101/2023.08.17.553594

**Authors:** P. D. Rich, S. Y. Thiberge, B. B. Scott, C. Guo, D. G. Tervo, C. D. Brody, A. Y. Karpova, N. D. Daw, D. W. Tank

**Affiliations:** Princeton Neuroscience Institute, Princeton University, Princeton, NJ, USA; Department of Psychological and Brain Sciences, Center for Systems Neuroscience, and Neurophotonics Center, Boston University, Boston, MA, USA; Janelia Research Campus and Howard Hughes Medical Institute, Ashburn, VA, USA; Princeton Neuroscience Institute and Howard Hughes Medical Institute, Princeton University, Princeton NJ, USA; Princeton Neuroscience Institute and Department of Psychology, Princeton University, Princeton, NJ, USA; Princeton Neuroscience Institute Princeton University, Princeton, NJ, USA

## Abstract

The precise neural mechanisms within the brain that contribute to the remarkable lifetime persistence of memory remain unknown. Existing techniques to record neurons in animals are either unsuitable for longitudinal recording from the same cells or make it difficult for animals to express their full naturalistic behavioral repertoire. We present a magnetic voluntary head-fixation system that provides stable optical access to the brain during complex behavior. Compared to previous systems that used mechanical restraint, there are no moving parts and animals can engage and disengage entirely at will. This system is failsafe, easy for animals to use and reliable enough to allow long-term experiments to be routinely performed. Together with a novel two-photon fluorescence collection scheme that increases two-photon signal and a transgenic rat line that stably expresses the calcium sensor GCaMP6f in dorsal CA1, we are able to track and record activity from the same hippocampal neurons, during behavior, over a large fraction of animals’ lives.

## Introduction

The hippocampus is crucial for episodic memory^1^, a hallmark of which is that events can be remembered over a whole lifetime^2^. The precise neural mechanisms within the hippocampus that contribute to this remarkable persistence remain unknown^3^. Monitoring the activity of neurons over the whole lifetime of animals during ethologically relevant behaviors is a crucial technical advance that is needed to fill this gap.

Head-fixed preparations are widely use in neuroscience research, providing optical access to the brain for cellular calcium imaging^4^ and optogenetic stimulation^5^ as well as the ability to present controlled stimuli to animals^6,7^. However, such preparations severely limit the complexity and naturalism of behavior that can be studied. The study of spatial navigation at naturalistic scales^8^ and in complex environments^9^, as well as episodic^10^ and social tasks^11^ requires the full response repertoire of a freely-moving animal. Rats can be trained to voluntarily head-fix^12^, offering the best of both worlds: stimulus control and brain access during head-fixation and the naturalism of freely-moving behavior otherwise.

Previous systems in rats^12,13^ and mice^14–17^ have relied on a mechanical clamp to restrain a head-plate attached to the head of the animal. Head restraint is unavoidably stressful^18,19^, and animals must be gradually habituated to it, for instance requiring progressive increases in the pressure of clamping pistons. The ability of animals to self-release is also critical^12,17^, but the mechanical solutions implemented to date impose a delay and are a potential point of failure; even one unsuccessful release attempt is enough for the system to become aversive for animals^17^.

We present a new approach that improves upon existing systems by eliminating any actual restraint of the animal. Instead, animals are trained to clip into a magnetic coupling from which they can leave at any point. Instead of being trained to tolerate restraint, animals just learn to hold still. This system is fundamentally less aversive and, since it employs no moving parts, eliminates the risk that failure of the system could derail long-term longitudinal studies.

Cellular resolution two-photon calcium imaging requires high stability^4^, and, for voluntary head-fixation, micron-scale repeatable registration between insertions. Kinematic clamps are widely used in mechanical engineering to repeatably reposition objects in space^20^; and when integrated into a voluntary head-fixation system allow *in vivo* two-photon calcium imaging^12,13^. We present a magnetic, full kinematic system with a geometry optimized for long-term voluntary use in rats. Ultra-hard and wear-resistant bearing surfaces provide the stability and reliability for routine two-photon imaging over long periods.

Due to the inherent optical sectioning of two-photon microscopy, every emitted photon from a sample is effectively signal^22^. Conventional two-photon microscopes use a single objective to both focus excitation light and collect emitted light; any emitted fluorescence that does not enter the front element of the objective with an appropriate angle is lost. Capturing these lost photons would improve the signal-to-noise ratio, increasing possible imaging depth and reducing the excitation power required (to prevent phototoxicity). Hybrid imaging/non-imaging objectives^23^, fiber light guides^24^ and parabolic epi-fluorescence reflectors^25,26^ have all been used to successfully improve collection efficiency, however, existing designs require the addition of complex collection optics to the microscope. We present a simple epi-fluorescence collection scheme that is based on a parabolic reflector placed close to the sample. A single drop-in piece replaces the cylindrical canula widely used for deep brain imaging^27^ and redirects otherwise lost photons into the existing collection path.

Genetically expressed calcium indicators in transgenic animals are the preferred method for neuronal population imaging in mice^28,29^, allowing large populations of neurons to be monitored over long periods^30^. Rats offer the opportunity to study more complex cognitive behaviors compared to mice but previous calcium imaging in rats has relied on the viral expression systems^12,13,31^ which limit the scope of experiments that can be performed. Transgenic rats have been developed^32^ that express GCaMP6f under the Thy-1 promotor and demonstrated for cortical imaging. We report a line generated using this technique that shows strong, stable and sparse expression in the CA1 region of the hippocampus, making it ideal for long-term population imaging of the same neurons.

We show that the magnetic head-fixation system achieves the stability and trial-to-trial reproducibility required for two-photon imaging; it is fast for animals to learn and animals are comfortable performing hundreds of trials over multiple sessions for months and years. Combined with stable transgenic expression of GCaMP6f in rats and the improved collection efficiencies of the epifluorescent collection cannula, we have been able to record from the same neurons in the hippocampus, during behavior, across the majority of animals’ lifetimes, opening up a domain of experiments with profound implications for the study of brain function.

## Results

### Design of the magnetic head-fixation system

The design of the magnetic head-fixation system was based on two constraints, micron-scale registration of the head plate between insertions and ease of use for the animal. Micron scale registration was achieved by kinematic design based on the Kelvin coupling^20^: an arrangement of a cone with three contact points, a vee-groove with two contact points, and a flat piece with a single contact point. These six contact points will non-redundantly constrain the six degrees of freedom of the head-plate according to the principle of exact constraint (Fig. 1A,B). We used tungsten carbide for the bearing surfaces of the cone and slot bearings, and hardened stainless steel for the flat bearings. Extreme hardness, wear resistance, and electrical conductivity make tungsten carbide the ideal material for this application, especially in the front bearings which are more subject to wear. Only following extended periods of continual use (many months, corresponding to tens of thousands of individual fixations) did we observe some wear and brinelling of the bearings, evidenced by intermittent failure of the electrical contact mechanism despite sufficient kinematic alignment. In these cases, replacement of the ball bearings or stage bearings resolved the issue.

**Figure 1.**
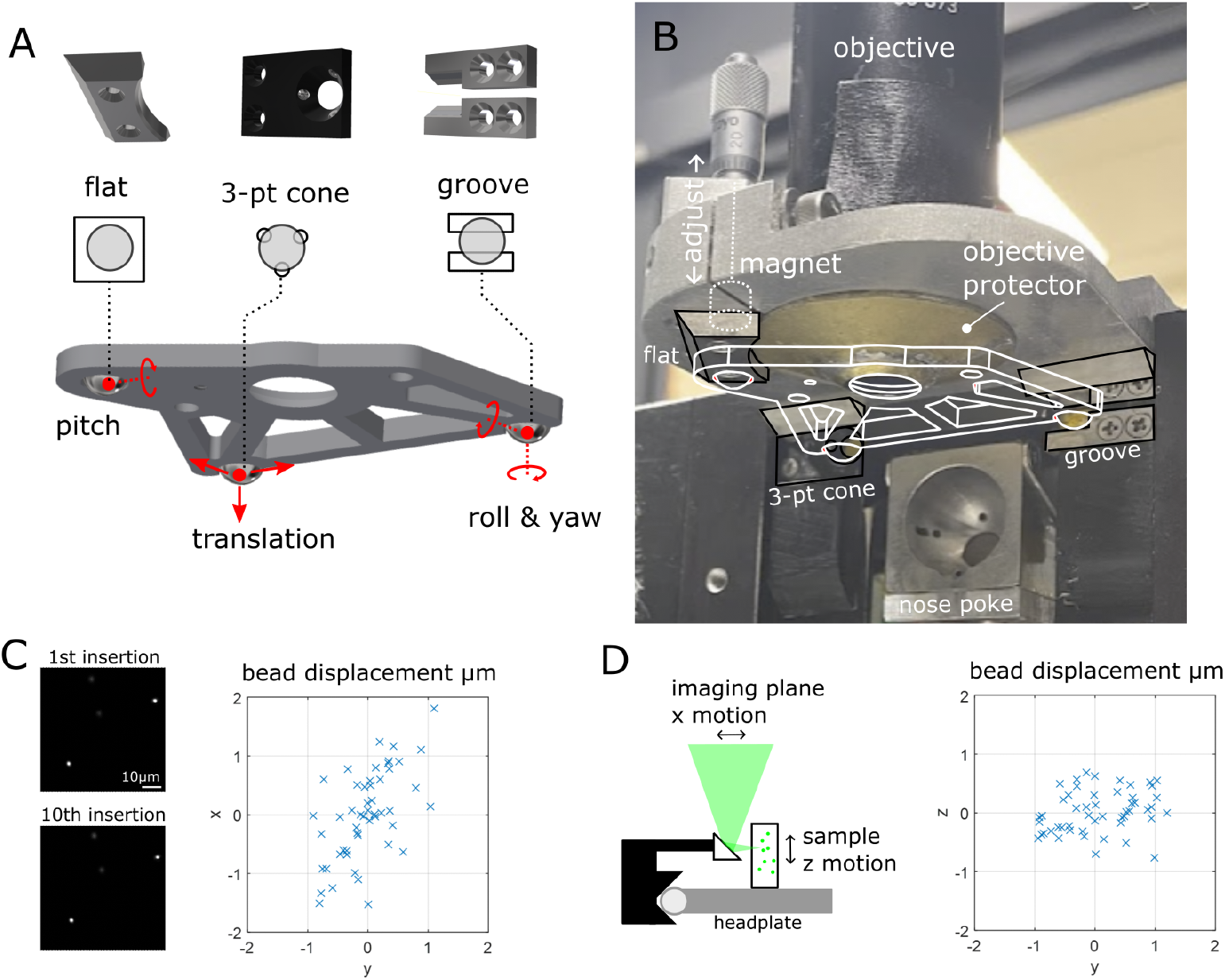
Magnetic voluntary head-fixation system. **A** The core of the system is the head plate with three ball bearings and three bearings of the Kelvin kinematic coupling. The 3-point cone, groove and flat bearings non-redundantly constrain the six degrees of freedom of the head plate. **B** Photograph of the bearing system showing interface with head plate (white, wireframe). The brass objective protector prevents the animal from touching the front aperture of the objective while not occluding the optical excitation or emission. The magnet and adjuster are shown for the flat bearing only, two other magnets and adjustment screws sit behind the cone and groove bearing. **C** First and tenth images of a fluorescent bead sample taken during *in vitro* testing of kinematic registration and the bead displacements in x and y for all insertions during the test. **D** A microprism attached to the bearing system allows the translation of sample z motion to imaging plane x motion. The bead displacements in x and z are shown for all insertions of the test.

Ease of use for the animal means that animals should be able to easily insert and exit at will from the system. The nose poke (Fig. 1 B) performed the initial coarse alignment of the head through alignment of the snout. Forward translation of the head, achieved by moving the nose poke back, is sufficient to seat the two front bearings which constrain the translation, roll and yaw of animal’s head. The final pitch degree of freedom is constrained by the flat bearing; adjusting the pitch of the nose poke during training allows the animal to readily learn the correct position.

We used magnets behind each bearing to provide a light attractive force on the ball bearings of the head-plate, both to aid the seating of the bearings and to help the animal keep the head-plate correctly seated during fixation. The system operates with no moving parts and animals could easy overcome the magnetic attraction and so break fixation at will. The magnetic force was started at zero initially and was increased by changing the distance from the magnet to the ball bearing with a micrometer screw. Animals were incentivized to complete fixations by delivering a chocolate milk reward at the end of the fixation duration.

### Registration is accurate *in vitro*

We assessed the reproducibility of the system without an animal by measuring the registration accuracy using a fluorescence bead sample (Fig. 1C). Across 50 insertions we measured the RMS of the displacement to be 0.81, 0.47, and 0.36μm in the x y and z dimensions respectively. This is in line with previous kinematic registration errors using ruby ball bearings and steel stage-bearings^21^.

### Reflective conical cannula for enhanced epifluorescence collection

Previous attempts to capture additional fluorescence in two-photon microscopy have utilized reflectors and relays to redirect light to collection optics^23,26^. In this system, we modify the widely-used canula prep for imaging deep brain structures such as the hippocampus^27^ to optimize light capture. By using a conical rather than cylindrical cannula, and polishing the interior wall to a mirror finish, we formed a collection optic that approximates the ideal parabola^25^ with a focal point at the imaging plane (Fig. 2A). Emitted light that would be otherwise lost is redirected up into the front element of the objective (Fig. 2B,C) and recorded with the existing collection optics. We measured the improvement in the optical collection of the polished conical cannula over the conventional cylindrical cannula and found up to a 1.5-fold increase in the collected fluorescence (Fig. 2D). The extra collection efficiently allows imaging of the CA1 pyramidal layer through the strongly scattering white matter of the alveus (Fig. 2E) which is thicker in rats compared to mice.

**Figure 2.**
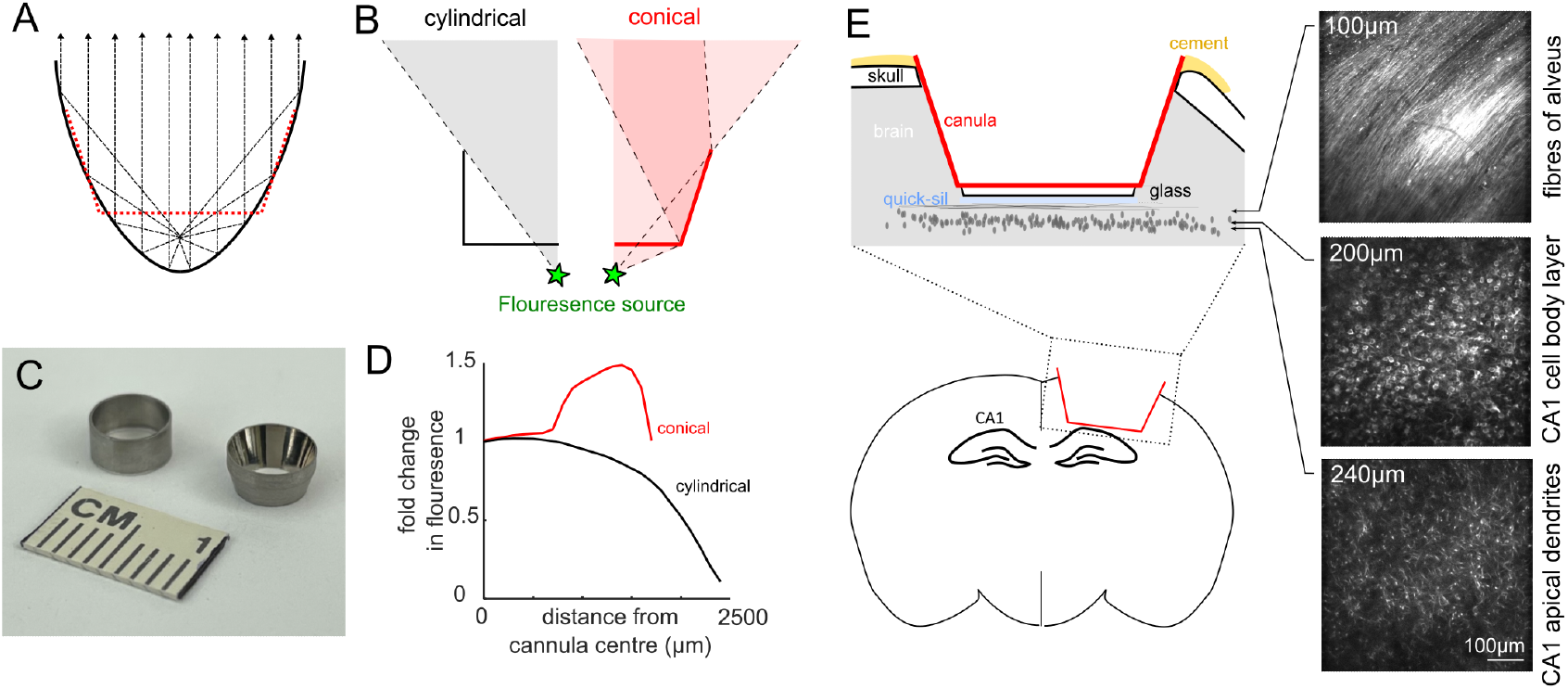
Conical cannula increases collected fluorescence and allows imaging of GCaMP6f expression in dorsal CA1. **A** A parabolic reflector redirects rays from a point source at its focus to a collimated beam. A conical cannula (red) approximates the parabolic shape, with the focus lying below the bottom of the canula at the approximate imaging plane of the CA1 pyramidal layer. **B** Compared to a standard cylindrical cannula, the conical cannula redirects emitted fluorescence that would otherwise be lost into the front aperture of the objective. **C** The inner surface of the conical cannula is polished to a mirror finish to improve collection efficiency. **D** Change in measured fluorescence at 150 μm below the cover glass (bottom of cannula). The conical cannula provides up to 1.5x increased fluorescence collection compared to a standard conical canula. The greatest improvement was measured at the edge of the canula, which corresponds to the location of the pseudo-focal point of the conical walls. **E** Installation of the conical canula over dorsal CA1. Schematic, *left*; two-photon images from an anaesthetized animal showing expression of GCaMP6f in the pyramidal cell layer at different depths bellow the bottom of the cover glass, *right*.

### Transgenic rats

Expression of genetically encoded calcium sensors has previously been achieved in rats using injection of viral vectors such as adeno associated viruses^12,13^. However, viral expression is difficult to maintain at healthy levels for long periods, severely limiting the scope of such experiments.

We used transgenic animals generated as previously described^32^, which carry the gene for GCaMP6f under control of the Thy-1 enhancer. Different rat lines have different levels of GCaMP6f in different brain region^29^. We selected a novel line, Thy1-GCaMP6f Line 8, that showed strong expression in the dorsal hippocampus (Fig. 2E, S1A). GCaMP6f was expressed in a sparse subset of hippocampal pyramidal cells in CA1 (Fig. 2E, S1B), and *in vivo* assessment of expression showed clear somatic localization (Fig. 2E).

### Animals are quick to learn magnetic head-fixation

Naive animals were trained to successfully align and hold their heads in the kinematic mount for at least 2 s in 988,203 (mean, sd) trials over 12.2,4.1 (mean, sd) sessions. Animals progressed rapidly through the three stages of training (Fig. 3), quickly learning to hold a nose poke and make contact with the front bearings (stage 1); hold contact with the front bearings and make contact at the rear bearing (stage 2); and finally hold contact with all bearings (stage 3). The magnetic attractive force for each bearing was introduced after animals began to make contact with each bearing, and was increased incrementally over a few sessions, to a level that made it easy for animals to sustain fixation whilst still being able to able break free (1.5 -2.5 N per bearing; overall measured head-plate pull out force <4N). The gradual introduction of the magnetic force meant that animals were not surprised by it, and we saw no evidence that it was aversive. This magnetic force both helped the alignment into the bearings and aided retainment as small forces generated by the animal did not cause it to lose fixation. Once proficient, animals typically performed daily sessions of 100s of fixations over multiple months and years, indicating a high degree of comfort with the apparatus (Fig. 3D).

**Figure 3.**
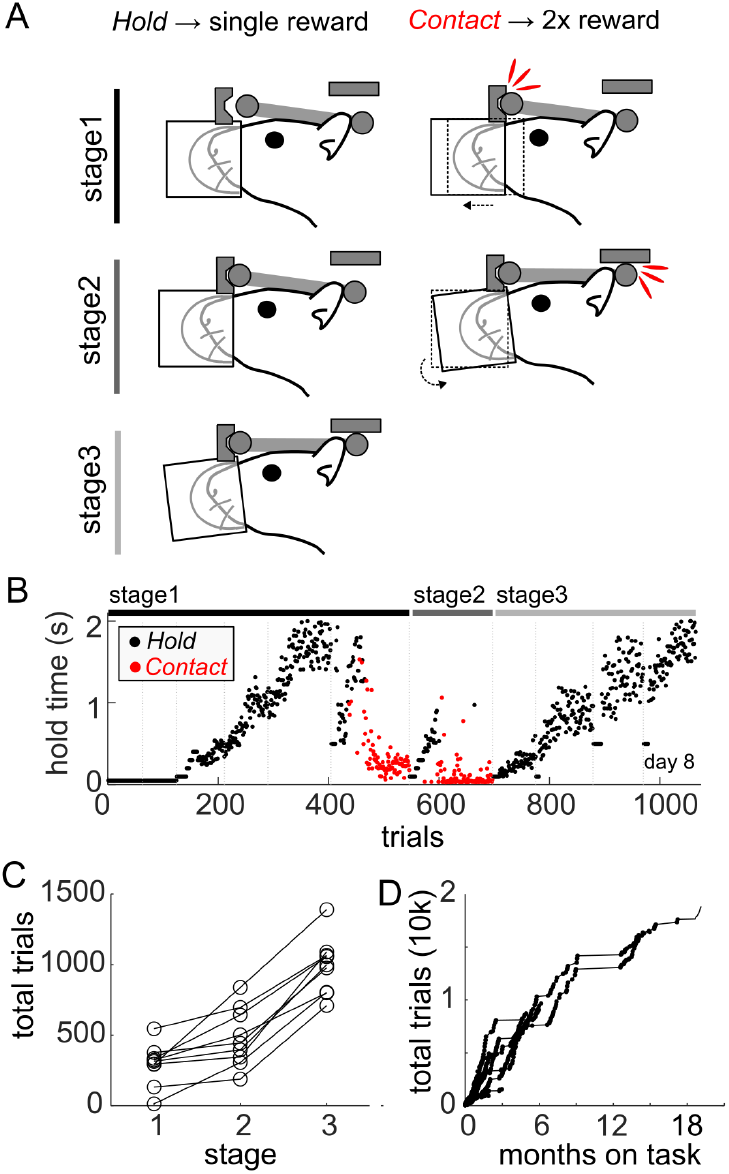
Behavioral training of voluntary magnetic head fixation. **A** Three stages of training. In the first stage animals receive a single reward for staying in the nose poke for the hold time; by increasing the required hold time, and moving the nose poke, the animal makes contact with the front bearings, which leads to an immediate double reward. Once animals are reliably making contact, they move to stage 2 which requires a hold on the front bearings; by increasing the required hold time, and rotating the nose poke, the animal makes contact with the rear bearing, which leads to an immediate double reward. Once animals reliably make rear contacts, they move to stage 3 which requires a hold on all bearings; hold duration is increased gradually. **B** Training profile for a single animal. Each point is a completed trial, either at the required hold time, or by making a “contact” in red. Vertical lines are separate sessions. **C** The total number of trials needed to complete each stage of training. **D** The cumulative number of trials completed over time. Once achieving holds on all bearings of at least 2s, animals are considered on task, and are able to continue performing indefinitely

### Long fixations

We targeted a fixation duration of 2 – 2.5 s to allow for a 1 s odorant stimulus presentation and a short delay period, which was readily achieved in all animals. Indeed, some animals remained clipped in to the system after reward delivery and during consumption of the reward despite no contingency for doing so, further indicating that the fixation was not aversive.

By increasing the duration of the delay period before the small reward was delivered, we were easily able to increase the fixations from 2.5 s to 4 s for an animal within a single session. In a separate experiment, we asked how long an animal would be able to remain head-fixed. We provided an animal with random intermittent rewards for as long as it remained head-fixed; it was able to remain head-fixed for up to one minute at a time (Fig. S2) demonstrating the level of comfort animals had with the system. This duration is comparable with previous reports in mice using a mechanical clamp and release switch^17^ which allows multiple self-contained trials to be presented per fixation insertion.

### Trial-to-trial registration and brain motion *in vivo*

Brain motion is a phenomenon of all *in vivo* imaging preparations and must be sufficiently small to be corrected by software. Image displacement, within trial (1.64, 1.19, 1.67 μm RMS in x,y,z axes, respectively) and between trials (which includes any kinematic registration error; 0.55, 0.38, 1.50 μm RMS) was similar to reported values for awake behaving rats^12^ and mice^4^. Motion within the imaging plane (x, y axes) could be corrected by existing non-linear motion correction algorithms (see Materials and Methods) leading to stable images (Svid_1). The influence of axial movement (z axes) cannot be corrected in software since the brain moves tangential to the imaging plane, however the range of displacement is modest compared to the axial point spread function of the microscope and the diameter of cell bodies. Close inspection of images did not reveal obvious differences in the view of fine neuronal processes which are most sensitive to axial motion (Fig. 4B, Svid_1).

**Figure 4.**
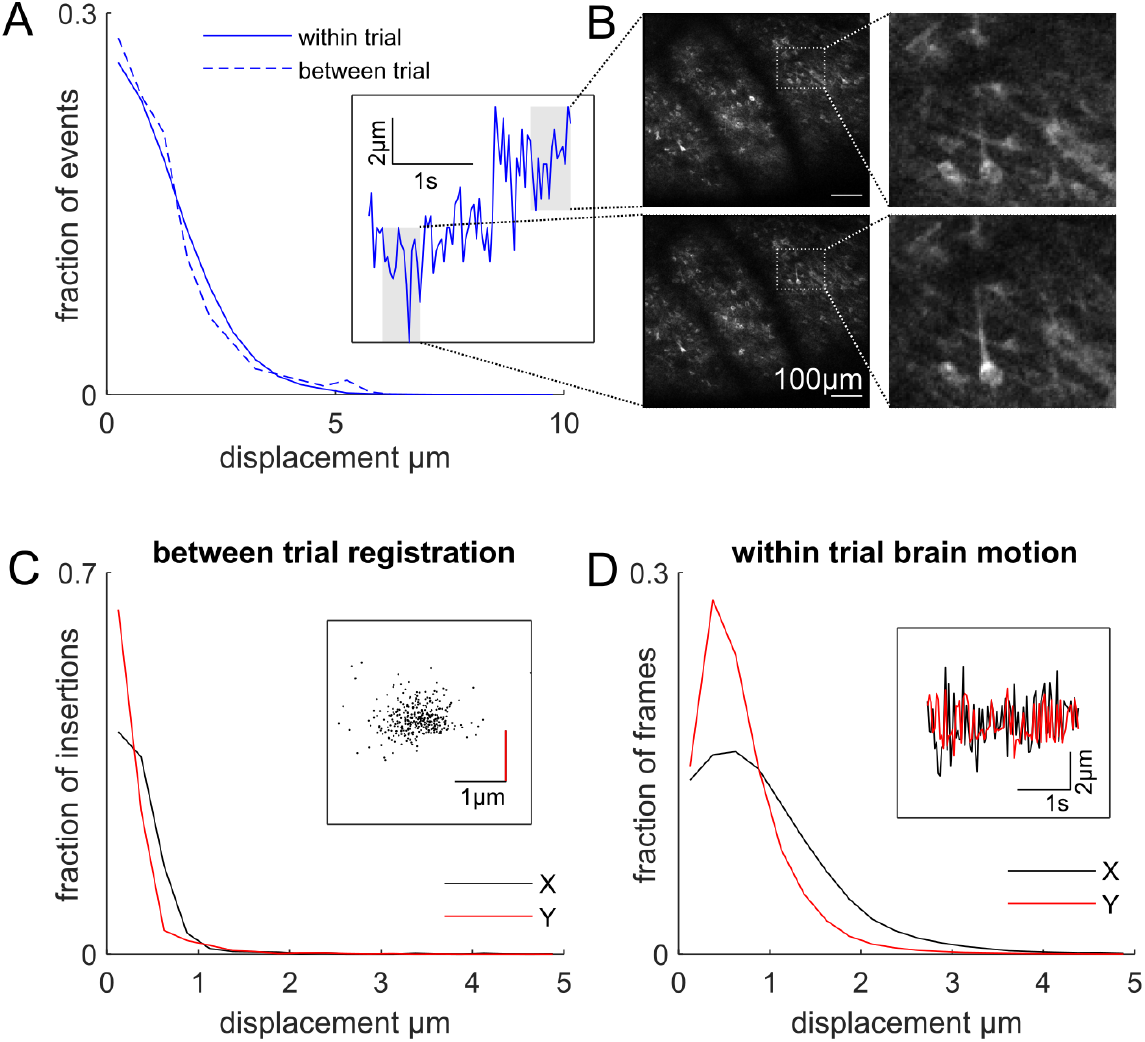
Trial-to-trial registration *in vivo*. **A** Histogram of the displacement of the brain in the z-axis, axial to the focal plane, for two rats. Inset shows the calculated frame-by-frame z-axis displacement for a single trial with a large range displacement. **B** Shows the average fluorescence fields of view for two 0.5 s periods at either end of the z-axis displacement range for the single trial inset in A. At the extreme range of such displacements, the same fine structures such as dendrites are visible in the image. **C** Histogram of the between trial registration displacements for six rats. Inset shows the spread of trial-to-trial displacements for a single session. **D** Histogram of the within-trial brain motion in x (anterior-posterior) and y (medio-lateral) for all rats. Inset shows the time course of the x and y displacements for a single trial.

### Calcium transients and odor responses in CA1

Over our large imaging fields of view, we were able to visualize the pyramidal cell body layer of CA1. As expected from the *in vitro* assessment, expression of GCaMP6f was sparse, with only a subset of cells being visible (Fig. 6B). We presented odorants to animals during fixation as part of an place-odor association task^33^: animals were trained that a large reward was available in a remote goal location depending on what odor was presented during fixation. We saw neurons in dorsal CA1 that responded selectively to different odors (Fig. 5A-D), and we were able to decode the presented odorant identity (Fig. 5E) at a population level.

**Figure 5.**
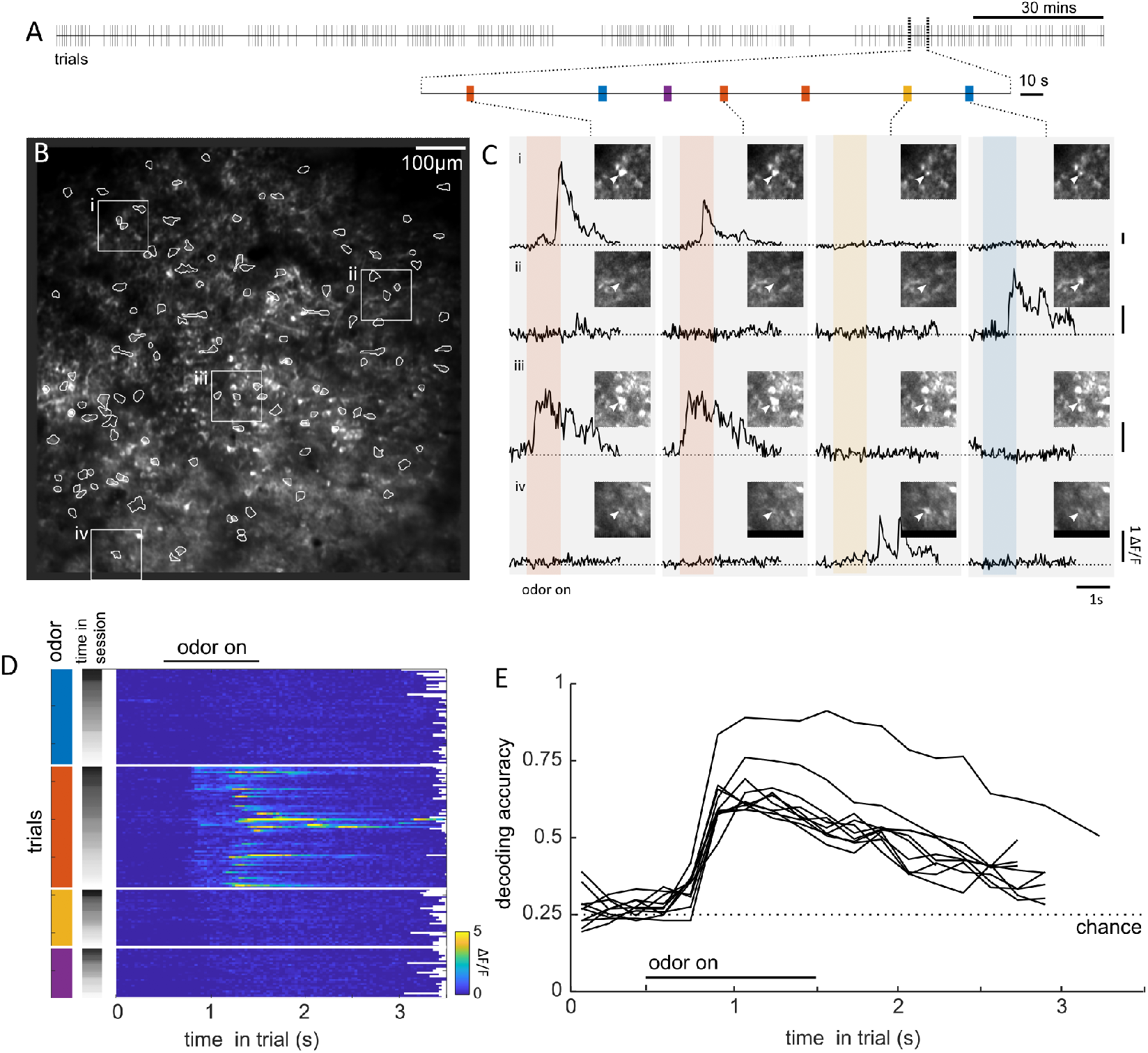
Odor responsive cells. **A** Time line of a behavioral session, with tick marks showing individual trials. **B** Average fluorescence field of view during the experiment. Active cells that were detected by the CNMF algorithm are circled. Numbered squared inserts (**i– iv**) correspond to regions centered on individual cells illustrated in C. **C** Fluorescence traces of four cells **(I – iv)** over four trials with various odors presented. Colored bar shows the 1 s period of odor presentation; different colors correspond to different odors. **D** Fluorescence traces for one cell (iii) for all trials in the session. Trials are first sorted by which odor was presented, and next by time in the session. **E** Odor decoding accuracy for a cross-validated linear classifier over the population of cells for 10 sessions for four rats. Dashed line shows the chance decoding level expected by chance.

**Figure 6.**
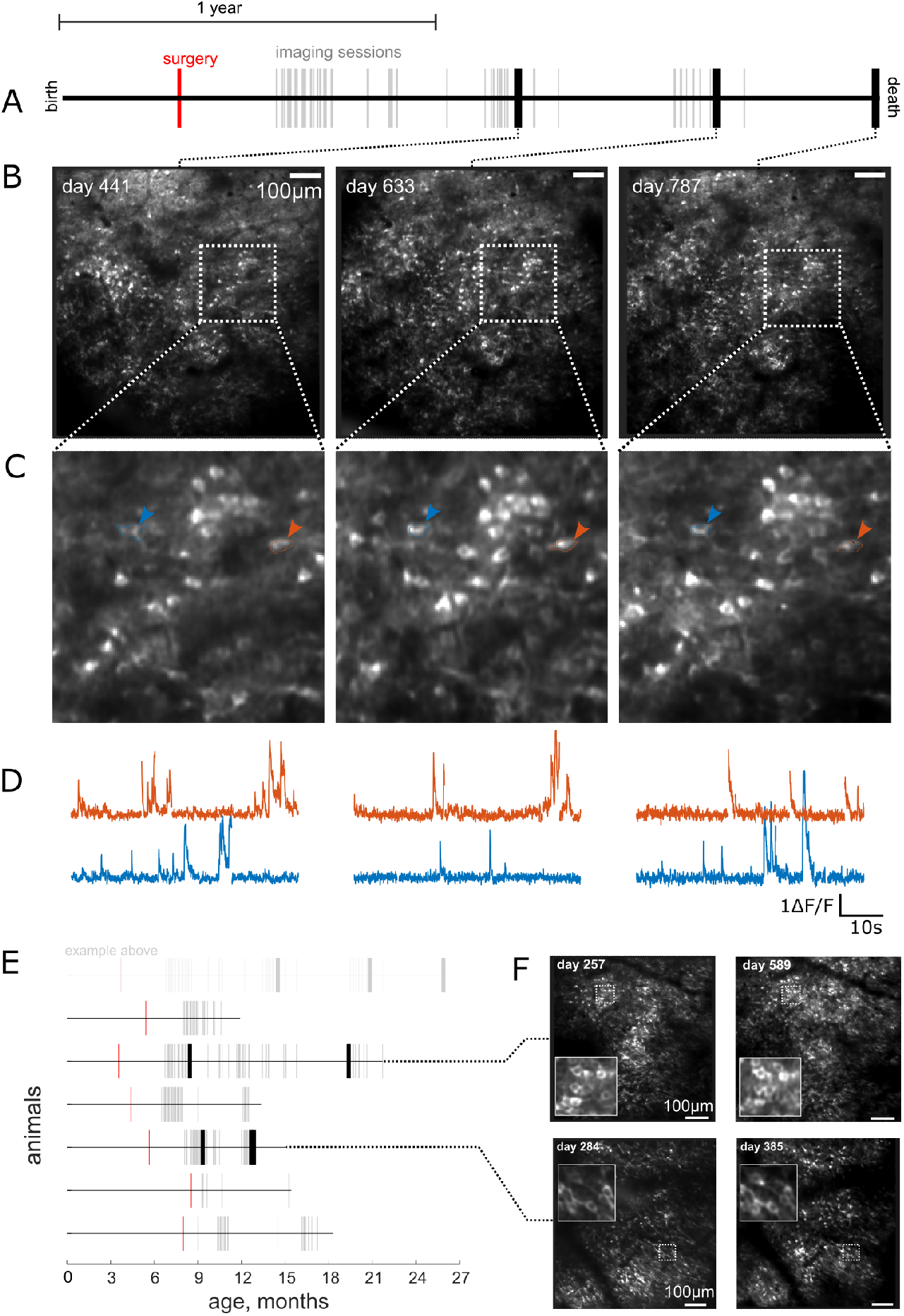
Lifetime imaging of the same cells. **A –** Life line of one rat; line shows the extent of the animal’s total life, 25 months. Awake behaving imaging sessions are shown as ticks. Bold ticks are example sessions illustrated below. **B** Average fluorescence fields of view for three example sessions. **C** Expanded views. Two corresponding cells are marked. **D** Calcium fluorescence traces for the two cells indicated above. Gaps in the calcium trace reflect the concatenation of individual trials. **E** Life lines of other animals, as in **A. F** Fluorescence fields of view and zoomed insert for two sessions for two animals.

### Long-term imaging

The expression of the calcium sensor and optical window clarity enabled imaging over a large fraction of animals’ lifespans. We were able to image hippocampal activity up to 21 months following surgery when one animal was 25 months old (Fig. 6A). Cells continued to show clear calcium transients (Fig. 6D), and the calcium indicator remained nuclearly excluded, indicating a level of GcaMP6f expression that does not affect cell function^28^. By returning to the same field of view, the same hippocampal cells could be tracked by comparing the morphology of individual neurons as well as the relative position of neighboring cells (Fig. 6B,C). We were able to track individual cells across a 19-month period in one animal (Fig. S3), other animals showed stability and longevity of the imaging window over many months (Fig. 6E,F).

## Discussion

Here we demonstrate three advancements that allowed us to track individual neurons over the entire adult life of animals as they perform a behavioral task: an entirely at-will magnetic head-fixation system that is safe and easy for animals to operate and is reliable over years of use, an epifluorescence collection canula that improves two-photon imaging signal, and a transgenic rat line that expresses GCaMP6f at stable levels in pyramidal cells of the hippocampus.

The magnetic voluntary head-fixation system described here is a substantial improvement over previous designs that have depended on mechanical restraint of animals^12,14,15,17^. The system provides the registration accuracy required for two-photon imaging through a kinematic clamp design^12,21^ using ultra-hard wear-resistance tungsten carbide and hardened stainless steel bearings. Since the system does not have any moving parts, it is also failsafe; there is no possibility of animals becoming trapped due to equipment failure. This allows the possibility of long-term, unsupervised, high throughput, home cage experiments with fewer safety and welfare concerns than existing systems. Adapting the system for use in mice would require a reduction in scale and the magnetic forces, and is feasible given the previous demonstration of kinematic clamps^21^ and voluntary head-restraint in this species separately^14,16,17^. Adapting this system for non-human primates offers a compelling direction for minimizing the invasiveness of these experiments, and could take advantage of the mature engineering principles required for designing high-load kinematics^20^. Although current head-mounted two-photon microscopes have begun to provide sufficient quality images to enable freely moving physiology^34^, they cannot approach the flexibility and extendibility of a benchtop system. Sophisticated techniques such as two-photon optogenetic stimulation^35^ and other exotic optical approaches can, at present, only be performed using head-fixed animals.

Building on previous approaches to improve epifluorescence collection in two-photon microscopy^26^ we were we were able to achieve a 1.5 fold increase in signal using an easy-to-fabricate cannula piece that approximates a parabolic reflector. We designed the current conical cannula shape for hippocampal imaging with a long working distance air objective but the current geometry would also be compatible with water immersion objectives commonly used for i*n vivo* two-photon microscopy, since they have a similar half angular aperture (37.0° for NA_water_= 0.8, 36.9° NA_air_=0.6) and sufficient working distances. By relaxing the current manufacturing constraints of straight cannula wall and performing end-to-end optical simulations, signal collection and field flatness could be further improved.

We add to the existing Thy-1 GCaMP6f rat lines previously reported^32^ with a line that shows strong, sparse and stable expression in pyramidal cells of the CA1 region. As well as being useful in our application of magnetic head-fixation, these current transgenic animals could be used to monitor calcium activity in freely moving rats using head-mounted two-photon microscopes^34,36^, head-mounted single photon microscopes^31,37^ or for fully head-fixed rat behavioral tasks^6^.

Tracking activity in the same cells over long periods is challenging. In electrode recordings, gliosis reduces signal^38^ and electrode movement changes the recorded waveforms of cells^39^. Reports of cells tracked over time have relied on stability of recorded waveforms^40^, stable tuning responses^41^, flexible electrodes^42^ or algorithmic post-hoc correction of electrode movement^43^. However, these methods are all indirect and not robust enough to enable routine lifetime recording, especially in the presence of representational drift^44^ or age-related changes in waveforms^42^ reported.

Calcium imaging provides a direct view of cells. Single photon imaging has been used to track cells over multiple days^37,45^, but is challenging over longer periods due to lack of optical sectioning, light scattering and its ability to only visualize active cells^46^. Using two-photon imaging, the morphologies of individual cells are clearly visible, which has enabled longitudinal structural imaging of dendritic spines *in vivo* for long periods^47,48^. Using genetically encoded calcium sensors the activity of the same neurons can recorded over months^49^, and as we demonstrate here, for well over a year. A sufficiently sparse and stable expression of the calcium sensor makes an unambiguous determination of cell identity across sessions relatively straightforward based on the unique constellation of cell positions. Future efforts to track cells across extended periods will be facilitated by having a similarly sparse labeling strategy as we have presented here.

The ability to follow the same cells over the lifetime of the animal has profound implications for the study of aging, and particularly, the study of age-related changes in the neural representations of memory. For example, continual optical access to the hippocampus and other brain regions over the lifetime of an animal would allow researchers to link the cellular and behavioral dimensions of Alzheimer’s disease to the *in vivo* measurement of amyloid plaques^51^. More generally, these techniques enables the long-term experiments that are necessary to link neural dynamics to the process of age-related cognitive decline^50^.

## Methods

### Animals and Surgery description

All procedures performed in this study were approved by the Institutional Animal Care and Use Committee at Princeton University and were performed in line with the Guide for the Care and Use of Laboratory Animals (National Research Council, 2011). We used 9 (5 male and 4 female) rats that expresses GCaMP6f under the thy-1 promotor aged between 3 and 9 months at the time of surgery. Animals were generated as previously described^32^, 7 animals were included in the long term imaging experiment following initial training on head-fixation.

Rats underwent analogous surgical procedures as previously described in mice^27^ to obtain optical access to the dorsal hippocampus. Surgery was performed under aseptic conditions, and animals were anesthetized with isoflurane (3.5% induction, 0.5 -1% for maintenance). Animals received pre-operative buprenorphine (0.02mg/kg) and either post-operative meloxicam (1mg/kg) or buprenorphine for analgesia. Body temperature was maintained with a heating pad (Harvard Apparatus). The skull was exposed and the periosteum was retracted. A craniotomy was performed on the right hemisphere over the dorsal CA1 region of the hippocampus (4.2AP, 3.0ML) The overlying cortex was aspirated to exposure the fibers of the external capsule, which was carefully retracted to exposure the fibers of the alveus. A thin layer of Kwik-Sil (WPI) was squeezed on the exposed area and the custom conical canula (see below) was quickly lowered and cemented in place with adhesive cement (C&B Metabond, Parkell). A titanium base head-plate (with screw holes to later accept the kinematic head-plate) was cemented to the skull. Following surgery animals were left for ∼1 month before the imaging window was assessed under anesthesia. At this point the kinematic head-plate (Fig. 1A) was installed into the base head-plate. The kinematic head plate was custom designed with 0.125” thick titanium using finite element analysis (Inventor, Autodesk) to optimize rigidity to weight. Three hardened stainless steel (440c, Mcmaster Carr) ball bearings were aligned to head-plate and a thin bead of conductive paint was applied to the joint to ensure electrical continuity across the head-plate. The ball bearings were then attached to the head-plate using the same adhesive cement used in the surgery.

### Conical cannula

The conical cannula was constructed from 420 series stainless steel chosen for its combination of corrosion resistance, magnetic (for holding the canula during implantation) and high surface finish achievable. The canula was 2.6mm in height with 4mm and 5mm diameter ends. The canula was turned on a precision lathe (Monarch 10ee), which allowed a high surface finish to be achieved on such a small piece. Following machining the interior of the cannula was polished with diamond polishing pins (Dedeco) and polishing paste (Simichrome). A 4mm diameter circular glass coverslip (Deckglaser) was glued to the bottom with UV curing glue (Norland 81).

### Imaging system

We performed imaging with a custom two-photon microscope controlled with the Scan Image software (Vidrio) running in Matlab (Mathworks). Laser illumination was provided by a Ti:Sapphire laser (Chameleon Vision II, Coherent) operating at 920 nm. We used a long working distance air immersion objective lens (20x/0.6NA, Edmunds optics) and a GaAsP photomultiplier tubes (H10770PA-40, Hamamatsu). The beam power was modulated by a Pockels cell (350-80 LA BIC -02 Conoptics) and the power used for imaging, measured at the front of the objective, was 150-200mW. Horizontal scans of the laser were achieved using a resonant galvanometer (Thorlabs). Typical fields of view measured approximately 600 × 600μm and data were acquired at 30 Hz.

### Bead measurements

1μm green fluorescent beads at a 1:1000 dilution were embedded in 3% agar gel for registration measurements. The agar solution was poured into a 5mm diameter canula with a glass cover slip glued to the bottom (Norland 91) and allowed to set. For x-y measurements the sample was attached to a head-plate which was inserted by the experimenter by hand into the bearing mount, 1s (30 frames) of data was taken. The head-plate was removed and reinserted by hand between each acquisition. For measurement of z-registration error we attached a prism with a reflective hypotenuse to the bearing system (Optosigma), and oriented the bead sample 90 degrees such that the glass cover slip bottom the cannula was oriented towards the prism^21^, Movement in the z-dimension would be translated into movement in the x-dimension.

### Fluorescence collection tests

To test the collection gain of the conical cannula we used a dilute fluorescein sample (0.1%) in a well with a cover slip glass over the top. Either the conical or cylindrical canula, without a glass bottom, were placed on the top of the clover glass. We recorded 100 frames from the center of each cannula to its internal edge in 100μm steps, and did the same sequence at different depths below the bottom of the cover glass. The average raw fluorescence values at each depth and distance from the canula center were normalized by the average fluorescence measured below the glass after removing the cannula, and scaled to the value at the center of the cylindrical canula for comparison.

### Magnetic at-will head fixation system

The kinematic magnetic head fixation system was designed to be structurally rigid, simple for animals to self-align, and the retainment of the head-plate within the bearings was facilitated with magnets. The three bearings surfaces were made of hard, electrically-conductive materials. The three-point cone uses three tungsten carbide ball bearings (0.094”, Mcmaster Carr) glued into an anodized aluminum bracket that arranged the balls in tetrahedral arrangement with respect the ball bearing in the head-plate; the groove bearing was made of 440c stainless steel with a press fit tungsten carbide insert (rectangular bar stock, Mcmaster Carr); the flat bearing was constructed of hardened 440c stainless steel. Behind each bearing was a neodymium magnet (K&J magnets) glued to micrometer screws (Mitotoya) which were used to finely and reproducibly adjust the strength of retainment of each ball bearing. A spring scale was used to calibrate the retainment force of the micrometer settings.

Contact with the each bearing and the ball bearing of the head plate was measured by a low voltage (15v at 68mA) electrical continuity circuit. The circuit detected which of the three bearings were seated corrected. If all three bearings were seated correctly, then a signal was sent to open the shutter on the imaging laser, begin image acquisition and update the behavioral system. A stainless-steel nose poke was mounted on a dual axis translation and single axis goniometer stage (OptoSigma), it allowed the coarse alignment of the animal’s head and positioned a lick tube at the correct position so animals could lick while still head fixed. An objective protector was made of thin brass sheet bent and soldered into a truncated cone, and protected the objective from the animal, whilst allowing free optical access to the cranial window.

### Behavioral training

Animals were food restricted to 85% of their free feeding weight prior to behavioral training, and maintained at this level during training. For some period when animals were not being trained they were given access to food ad libitum and returned to food restriction prior to training.

Stage 1 - Animals were first trained to poke at the stainless-steel nose poke to receive a chocolate milk drop reward (1 drop, Ensure milk chocolate flavor nutritional shake). Alternation with a nose poke at the rear of the chamber where they also received a chocolate milk drop. The length of time the animal had to hold its nose in the nose poke was slowing increased; the hold time was drawn from a distribution of times the bounds of which were increased during the session. If the animal was successfully making holds, the lower bound was increased 0.1s every 10 trials and the upper bound 0.2s every 10 trials; for a given session, this increase was stopped when the lower bound reached the value of start of session upper bound. During the hold period an audible frequency modulated tone was played to indicate to the animal an ongoing successful hold. The nose poke position was adjusted so that the head plate ball bearings would contact the two front bearings which triggered a larger immediate reward (2 drops).

Stage 2 - Contact with the front bearings now initiated the hold period which, as before was drawn from a range which was increased throughout a session. Contact with the rear, flat bearing now triggers the immediate large reward. In this condition the nose poke may be needed to adjusted in the pitch axes using a goniometer stage to enable contact with rear flat bearing. The holding force of the magnetic front bearings was increased in this stage, beginning at 0N it was increased up to 2.5N to aid the animal to maintain the hold, and was increased in small increments every 10-20 trials over the course of 1-2 sessions

Stage 3 - Contact with all bearings now initiated the hold period which, as before was drawn from a range which was increased throughout a session. Animals were required to maintain fixation for the hold duration to receive a reward. The magnetic attraction for the rear bearing was adjusted at this stage from 0N to 2.5N and again was increased in small increments every 10-20 trials over the course of 1-2 sessions, the lower bound of the hold duration was increased so that the hold period was fixed at a set value of 2-2.5s.

If animals did not complete a fixation (i.e. they did not remain fixated for the required hold time or make a contact to get a large reward during stages 1 and 2), there was a 1s time-out period before a new trial could be initiated. The illumination of a blue LED light inside the nose poke indicated to the animal when a new trial could be initiated.

### Odor guided navigation task

Once animals had learned to reliably clip-in to the head fixation system they were trained to perform an odor guided navigation task. Animals were trained that different odors corresponded to different reward location in a maze constructed of enclosed linear track elements. We delivered odors (acetic acid 20%, 2-propanol, propyl acetate, anethole, trans-cinnamaldehyde, nutmeg, cardamom, clove, rosemary, star anise) to animals using a custom built olfactometer. Air was directed to flow though vials containing odorants using PTFE valves (Neptune Research) at a flow rate of 0.5L/min. At the start of head fixation, blank air was delivered for 0.5s, followed by 1s of odorized air, followed by a scavenger vacuum through the system to evacuate any residual odorant for the next trial.

### Motion correction

Motion correction was performed as described previously^52,53^, imaging data were corrected for non-rigid brain motion using custom written MATLAB code based on a similar algorithm as NoRMCorre^54^ in which the image is divided into overlapping patches and a rigid translation is estimated for each patch and frame by aligning against a template. The maximum displacement out of all patches in a frame was used to calculate the displacement values for the x and y axes.

### Z-motion estimation

Motion perpendicular to the imaging plane (z-motion) was estimated by comparing imaging frames to a reference z-stack collected during anesthetized imaging session in two animals. The reference Z-stack was collected at the same coordinates as awake imaging and each image was made as the average of 100 frames taken at 1um spacing. In order to account for the non-ridged motion correction patches of the average motion corrected movie were correlated with corresponding patches in the reference Z-stack. The Z-motion for each frame was calculated as average frame of peak correlation for each patch.

### Roi extraction

Fluorescence traces corresponding to individual cells were extracted from the motion corrected images using Constrained non-negative factorization algorithm (CNMF)^55^. Initialization of the spatial components for CNMF was done as previously published as was classification of identified components into cell-like and non-cell-like categories^52,56^. All components were manually inspected and re-classified if needed. ΔF/F for each cell was calculated using the modal value of fluorescence in 3-min long windows as baseline fluorescence. Since CNMF only extracts cells that have calcium activity during imaging, silent cells that did not show activity during imaging were not included.

### Odor Classifier

We used a linear classifier approach to decode the odor identity. We used K-fold cross validation and trained a linear decoder on the population activity for cells that exceeded an average activity threshold of 0.03 ΔF/F across the whole session.

### Histology

Histology was performed to assess the raw GCamps6f signal^29^. Animals were anesthetized with an overdose of Ketamine Xylene and transcardially perfused with PBS and 4% Paraformaldehyde. Brains were extracted, and left in fixative for 12hrs and then transferred to 30% sucrose in PBS until the brain had sunk. Brains were sectioned on a freezing microtome (Leica) in 50μm sections and floated onto slides from a solution of DPBS +calcium. Slices were mounted on slides with prolong diamond +DAPI (Thermofisher) and left refrigerated for at least 12hrs before imaging. Slides were imaged with an epifluorescence microscope for whole brain expression pattern and a confocal microscope for cellular resolution images.

## Supporting information

Svid_1

Svid_2

## Author Contributions

P.D.R. performed the experiments and data analysis,

P.D.R., S.Y.T. and D.W.T. designed the experimental setup,

B.B.S., C.G., D.G.T., C.D.B., A.Y.K. Generated and screened the transgenic animal line,

P.D.R. wrote the manuscript with input from D.W.T.,

N.D.D., and D.W.T. supervised the project.

## Acknowledgements

We thank A. Song for assistance with the microscope, J. Gauthier for help with hippocampal surgeries and S. Koay for cell extraction software and assistance. We thank S. Lowe for assistance machining and H. Payne for comments on the manuscript.

## Supplementary videos

**Svid_1**

High magnification field of view for three trials (three columns) in the long fixation experiment. The top element is the raw calcium fluorescence movie, and below that is the same video but motion corrected.

**Svid_2**

Showing head-fixation and imaging during two trials of the odor guided navigation task. The animal head fixes (upper left panel) and imaging starts, scattered infrared light is visible from the un-shuttered imaging laser. During imaging the raw calcium imaging movie of a wide view of CA1 pyramidal cells is shown on the right. The colored square indicates the odor delivery duration. Calcium transients are visible in some cells. After the fixation period has been completed, choice doors open to the left and right, giving the animal access to the rest of the maze. The animal then runs to the correct remote location (bottom left panel) to collect the reward.

## Supplementary figures

**Fig S1.**
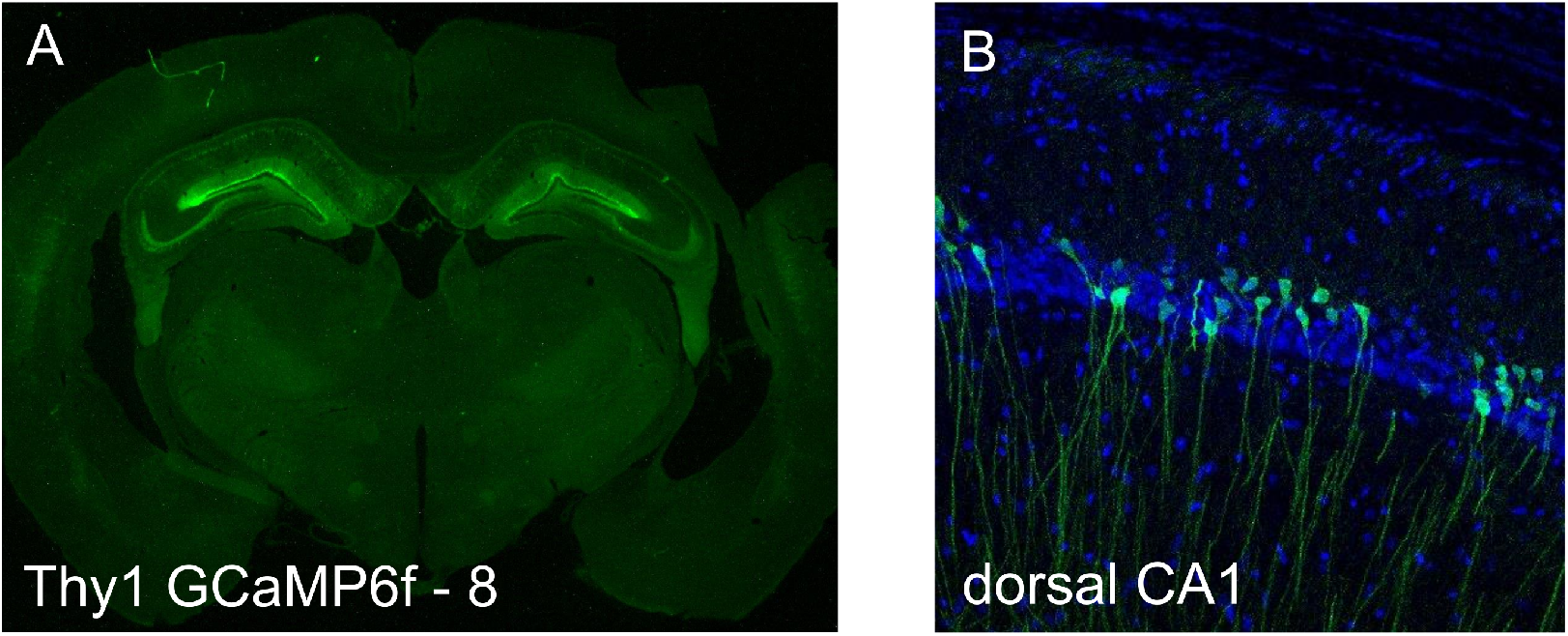
Transgenic expression of GCaMP6f. **A –** Epifluorescence microscopy image of native GCaMP6f expression (green) in a coronal section from a Thy1-Gcamp6f-8 rat. **B** Confocal microscopy image of the of native GCaMP6f expression (green) and DAPI (blue) of the CA1 layer of the dorsal hippocampus.

**Figure S2.**
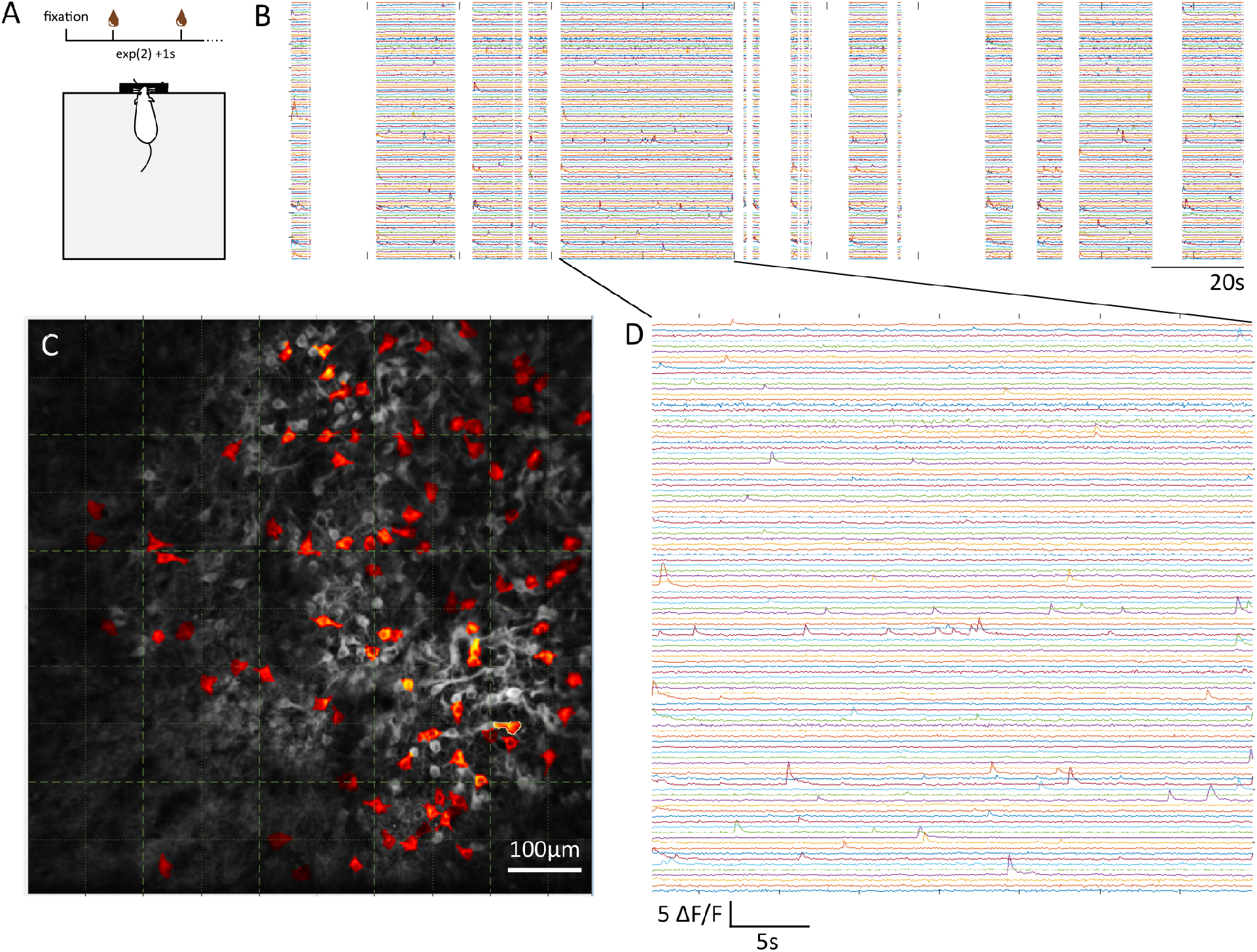
Long fixation imaging. **A –** Showing the long fixation experiment in which the animal is required to hold fixation for as an indefinite period, during which time it gets random rewards with an interval given by a random draw from an exponential with a mean of 2s plus 1s. **B** Calcium fluorescence time courses of active cells during fixation. **C** Field of view showing average fluorescence and highlighted active cells. **D** Zoom of calcium activity for a single fixation of 37s.

**Figure S3.**
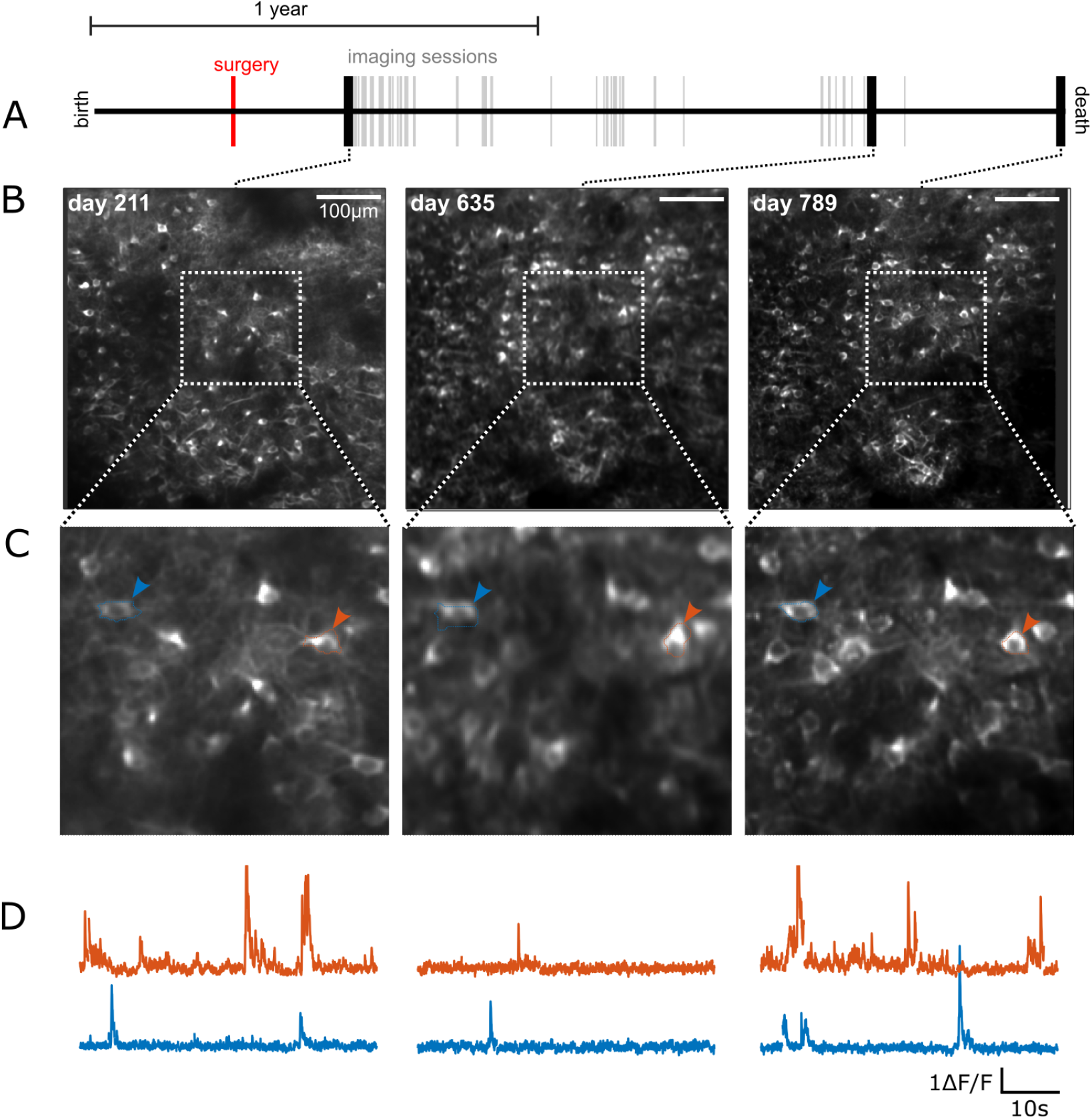
Same cells are visible and show activity over 19 months. Different field of view for the same animal shown in Fig. 6. **A –** Life line shows the extent of the animal’s total life, 25 months. Awake behaving imaging sessions are shown as ticks. Bold ticks are example sessions illustrated below. **B** Average fluorescence fields of view for three example sessions. **C** Zoomed in details of the fields of view. Two corresponding cells are marked. **D** Calcium fluorescence traces for the two cells indication above. Gaps in the calcium trace reflect the concatenation of individual trials. **E** Life lines of other animals, as in **A. F** fluorescence fields of view and zoomed insert for two sessions for two animals.

